# Uncovering patterns of atomic interactions in static and dynamic structures of proteins

**DOI:** 10.1101/840694

**Authors:** A. J. Venkatakrishnan, Rasmus Fonseca, Anthony K. Ma, Scott A. Hollingsworth, Augustine Chemparathy, Daniel Hilger, Albert J. Kooistra, Ramin Ahmari, M. Madan Babu, Brian K. Kobilka, Ron O. Dror

## Abstract

The number of structures and molecular dynamics simulations of proteins is exploding owing to dramatic advances in cryo-electron microscopy, crystallography, and computing. One of the most powerful ways to analyze structural information involves comparisons of interatomic interactions across different structures or simulations of the same protein or related proteins from the same family (*e.g.* different GPCRs). Such comparative analyses are of interest to a wide range of researchers but currently prove challenging for all but a few. To facilitate comparative structural analyses, we have developed tools for (i) rapidly computing and comparing interatomic interactions and (ii) interactively visualizing interactions to enable structure-based interpretations. Using these tools, we have developed the Contact Comparison Atlas, a web-based resource for the comparative analysis of interactions in structures and simulations of proteins. Using the Contact Comparison Atlas and our tools, we have identified patterns of interactions with functional implications in structures of G-protein-coupled receptors, G proteins and kinases and in the dynamics of muscarinic receptors. The Contact Comparison Atlas can be used to enable structure modeling, drug discovery, protein engineering, and the prediction of disease-associated mutations.

Contact Comparison Atlas website: https://getcontacts.github.io/atlas/

## Introduction

Advances in structural biology techniques such as crystallography, NMR, and cryo-EM have been propelling a rapid growth of protein structures from diverse protein families. Nearly 150,000 experimental structures are available publically (wwPDB consortium 2018). On top of that, the dynamics of these structures are being investigated by biomolecular simulation far more often owing to the development of faster and cheaper computers, leading to an explosion of data about structural dynamics (Hollingsworth and Dror 2018). This availability of unprecedented amounts of structural information has vast potential to be exploited for many discoveries and applications, but the sheer scale of this data brings new analytical challenges.

A fundamental way to analyze protein structures and simulations is through comparisons, and such comparative analyses can be informative in many different scenarios (Figure 1): (i) ligand binding affinity comparison: compare structures/simulations of the same protein bound to different ligands, or with and without a ligand, (ii) subtype selectivity: compare structures/simulations of related proteins bound to the same ligand, (iii) allostery and conformational change: compare structures/simulations of the same protein in different conformational states, and (iv) molecular evolution: compare structures/simulations of the same protein from different species.

**Fig. 1.**
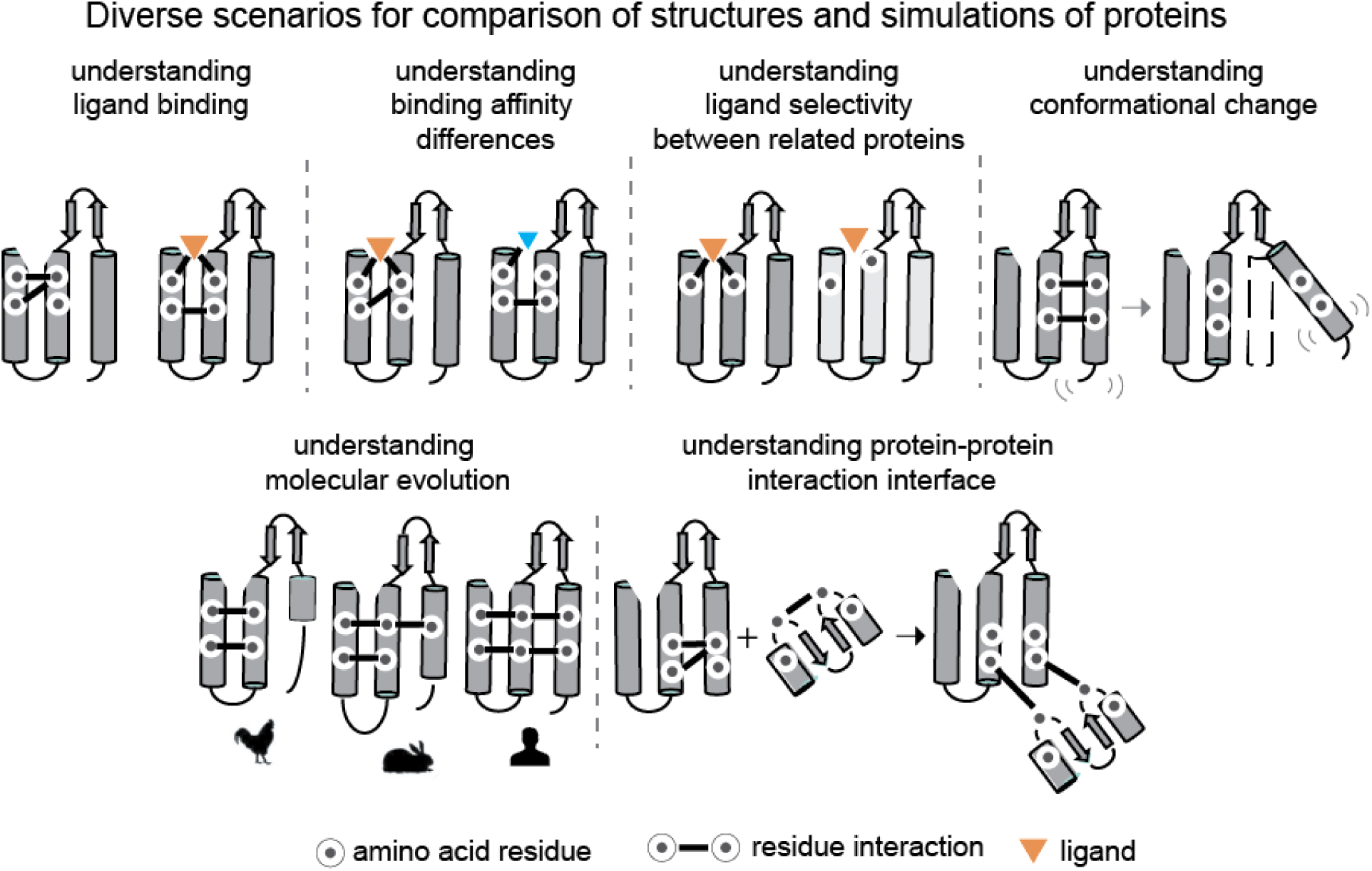
Different scenarios for comparative analysis of protein structures and simulations. (i) understanding ligand-binding: compare structures/simulations of same protein bound to different ligands, or with and without a ligand, (ii) understanding subtype selectivity: compare structures/simulations of related proteins bound to the same ligand, (iii) understanding allostery and activation: compare structures/simulations of the same protein in different conformational states, (iv) understanding molecular evolution: compare structures/simulations of the same protein from different species, (v) understanding protein-protein interaction interfaces: compare unbound and bound structures/simulations of interacting proteins, (vi) understanding protein nucleic acid interaction interfaces: compare unbound and bound structures/simulations of interacting proteins.

In such comparative analyses of protein structures and simulations, focusing on patterns of atomic interactions (*e.g.* hydrogen bonds) enables us to gain an in-depth understanding of function (Kayikci et al. 2018)(Doncheva et al. 2011)(Zhang, Perica, and Teichmann 2013)(Vass, Podlewska, et al. 2018)(Vass, Kooistra, et al. 2018). For instance, studying patterns of interactions can help identify amino acid residues important for ligand-binding (Venkatakrishnan et al. 2013) (Ngo et al. 2017), understand similarities in activation mechanisms (Venkatakrishnan et al. 2016) and understand specificity in protein interactions (Flock et al. 2015).

While comparison of interactions across multiple structures and simulations is informative, performing such comparisons can be cumbersome. This is due to the need to compute the interactions for each structure/simulation, map the structural correspondences between different proteins, compare the interatomic interactions, and finally visualize the similarities and differences in the interactions in a structurally intuitive manner.

Here, we present GetContacts – a software package to rapidly compute and compare interactions in both protein structures and simulations (Figure 2a). We have also developed interactive visualization tools that depict interaction patterns using both custom-made plots and structural renderings of proteins (Figure 2b). Finally, using these tools, we have built the Contact Comparison Atlas, a web-based platform for the analysis of interactions and interaction patterns in protein structures and simulations (Figure 3). We illustrate the use of GetContacts and the Contact Comparison Atlas to uncover shared and distinct functional properties in three biomedically important protein families: G protein–coupled receptors (GPCRs), G proteins, and kinases.

**Fig. 2.**
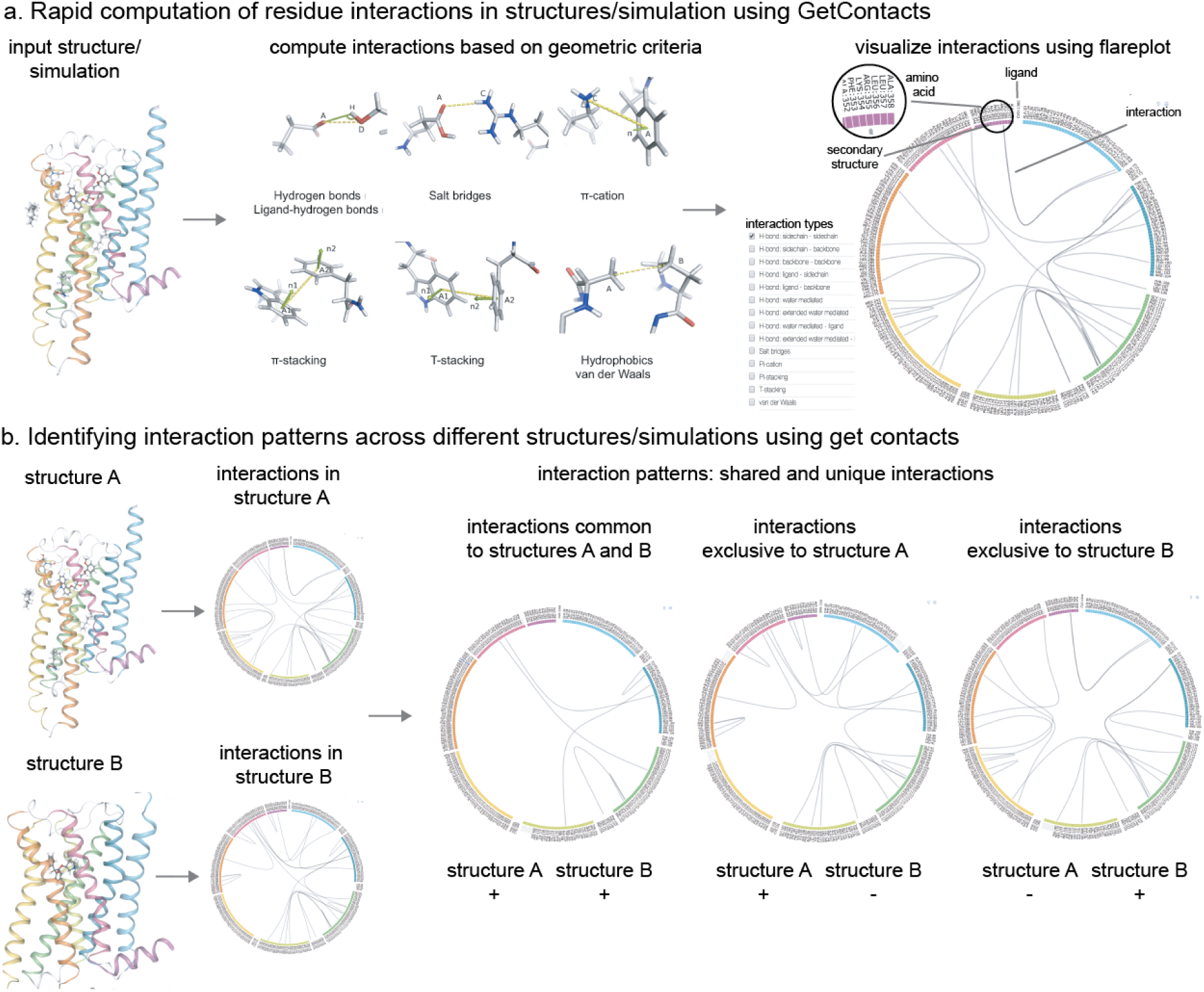
Computation and comparison of interactions using GetContacts. (a) Computation of residue contacts in structures and simulations using GetContacts. For a given structure/simulation, interactions and interaction frequencies are computed using distance and angle criteria. These interactions are visualized in 2D using flareplots and 3D using structure-based rendering. (b) Identifying interaction patterns across different structures/simulations using GetContacts. Once the interactions/interaction frequencies are computed for each structure/simulation being compared, shared and unique interactions are identified.

**Fig. 3.**
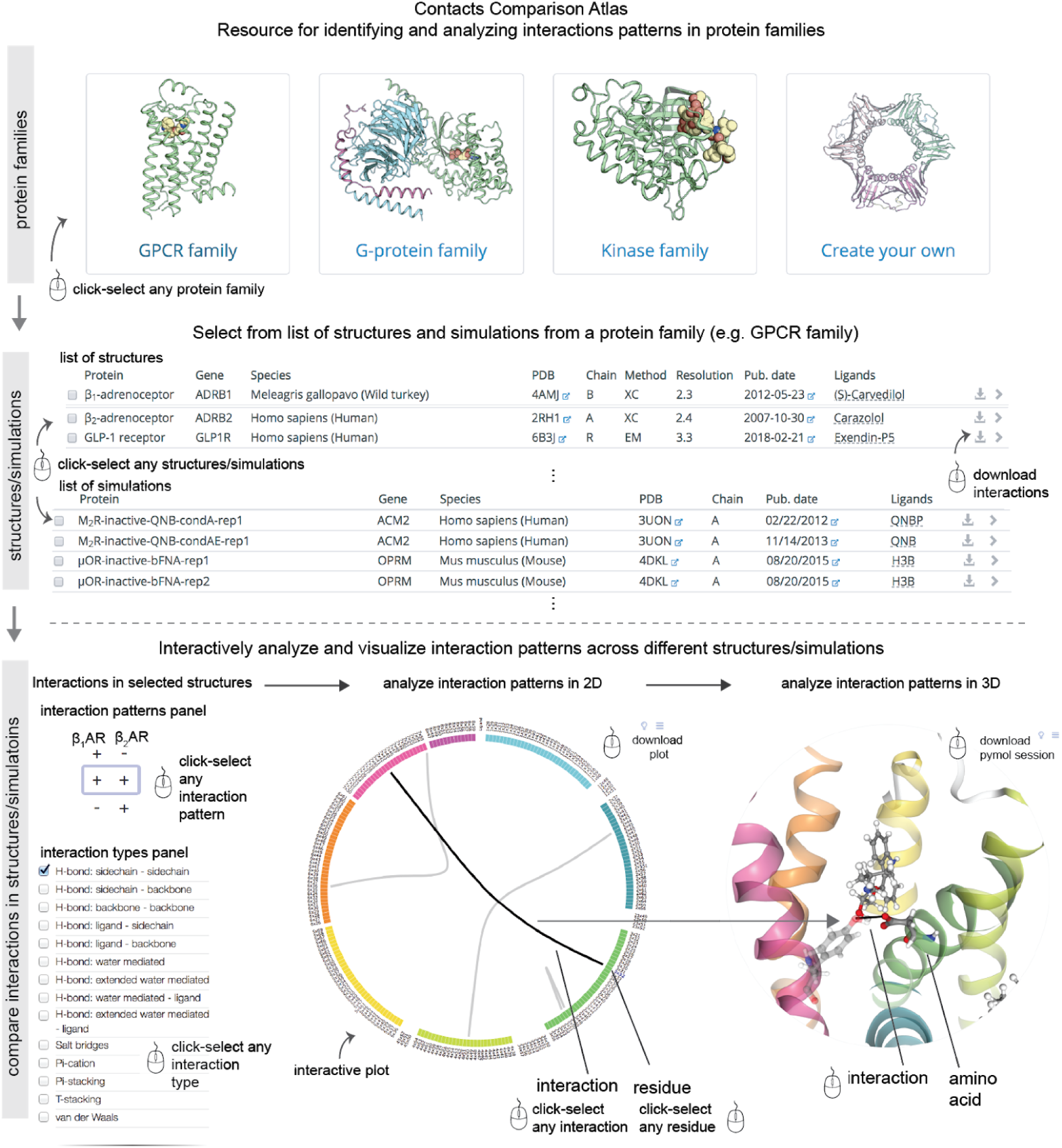
Framework of the Contact Comparison Atlas web-resource. **(top)** Selection of structures/simulations can be made from pre-compiled protein families (GPCRs, G proteins, kinases) or users can upload their own data to be selected from. **(middle)** Upon choosing the protein family/uploading data, the user is presented with a list of all the entries for selection of structures/simulations to be compared. (**bottom**) The comparison of interactions is presented in a new webpage for interactive analysis and visualisation. The interactive layout consists of four panels: interaction patterns panel, interaction types panel, flareplot panel and structure rendering panel.

## Interactive analysis of interatomic interaction patterns in proteins

Here, we describe the comparative analysis of a given a set of structures and/or simulations of proteins in three steps: (i) computing interactions, (ii) identifying interaction patterns (*i.e.* interactions that are shared across structures/simulations or that differ between them), and (iii) visualizing the results.

### Computing residue interactions from structures/simulations

We have developed a software package GetContacts for rapidly computing atomic interactions in protein structures and simulations. The computed interactions include hydrogen bonds, salt bridges, aromatic interactions (*i.e.* pi-pi stacking or pi-T stacking interactions), cation-pi interactions, and van der Waals contacts. GetContacts uses interatomic distances and angle-based criteria (see **Methods**) to identify interactions between pairs of amino acid residues and between amino acid residues and any bound ligands. For simulations, GetContacts identifies contacts present in each frame of the trajectory. GetContact’s high computational efficiency is important in this case, because individual simulation trajectories often contain many thousands of frames.

### Identifying interaction patterns across structures/simulations of the same protein in different conditions

In addition to computing atomic interactions, GetContacts can also identify interaction patterns (shared and unique interactions) across structures/simulations of the same protein in different conditions (*e.g.* different conformational states). Let us consider the example of comparing a pair of structures of the same protein but in two different conformational states: inactive state and active state. Once the interactions are computed for structures representing the two states (inactive and active), GetContacts identifies interaction patterns by separating the shared interactions and unique interactions into different sets: (i) interactions shared across both states, in the sense that the same amino acid residue pairs interact in both states, (ii) interactions unique to the active state, and (iii) interactions unique to the inactive state. While the number of possible interaction patterns is trivial when comparing two structures, the number of possible patterns grows exponentially with the number of structures being compared (2^n^; n=number of structures). GetContacts systematically identifies all interaction patterns irrespective of the number of structures.

In addition to identifying interaction patterns across multiple structures, GetContacts can also identify interaction patterns across simulations. GetContacts computes interaction frequencies at the residue level for each simulation, binarizes each potential interaction as present or absent based on a user-defined threshold for interaction frequency, and identifies shared and unique interactions across the simulations.

### Identifying interaction patterns across structures/simulations of different but related proteins

Comparing interactions across different proteins that are structurally related (*e.g.* different GPCRs) can be cumbersome because of differences in the amino acid sequences of the proteins being compared. We have solved this problem by taking advantage of protein-independent numbering schemes for amino acid residues that are based on structural alignments. Such numbering schemes include the Ballesteros-Weinstein numbers (Ballesteros and Weinstein 1995) or GPCRdb numbers (Isberg et al. 2014) for GPCRs, CGN numbers for G proteins (Flock et al. 2015), KLIFs-based numbers for kinases (Kooistra et al. 2016). For protein families where established numbering schemes don’t exist, structure-based sequence alignments generated by tools such as CCP4’s Gesamt (Winn et al. 2011) can be used to map structurally equivalent residues between proteins. Once the equivalent residues are mapped, using either numbering schemes or structure-based alignments, GetContacts can be used to identify patterns of atomic interactions.

Let us consider the example of comparing interactions between two different proteins belonging to the same family. Once the interactions are computed and structurally equivalent residues are mapped between the proteins, GetContacts identifies interaction patterns by grouping separately the shared interactions and unique interactions: (i) interactions present in both proteins, (ii) interactions present in the first protein only, and (iii) interactions present in the second protein only. In comparisons of more than two structures or simulations, GetContacts exhaustively identifies sets of interactions that are shared or distinct.

### Visual analysis of interaction patterns from structures and simulations

Once the interaction patterns are identified across multiple structures/simulations, interpreting the functional relevance of the patterns requires effective means to analyze and visualize the interactions. To this end, we have developed a web-based interface comprising four interactive panels (Figure 3): (i) an interaction-types panel used to specify which types of interactions (*e.g.* hydrogen bonds or van der Waals contacts) will be displayed, (ii) an interaction-fingerprint panel used to specify whether the displayed interactions should be present or absent in each structure/simulation, (iii) a 2D visualization of interaction patterns using ‘flareplots’, and (iv) a visualization of interaction patterns using 3D structural rendering. In the interaction-types panel, the different types of interactions are available in a drop-down menu. In the interaction-fingerprint panel, all the underlying patterns of interactions across the structures/simulations being compared are listed exhaustively. Each interaction fingerprint is a sequence of ‘+’ and ‘–’ signs corresponding to the column of structures/simulations being compared. A ‘+’ sign for a structure indicates that displayed interactions are present in that structure; for a simulation, it indicates that displayed interactions are present in that simulation at a frequency greater than or equal to a user-defined threshold. Conversely, a ‘–’ sign for a structures indicates the absence of an equivalent interaction in that structure; for a simulation, it indicates either that an equivalent interaction is formed at a frequency below the threshold or that an equivalent interaction is absent. For the 2D visualisation of the interaction patterns, we have developed a tailor-made visualization of interactions called a ‘flareplot.’ In a flareplot, the amino acid sequence is placed in a circular layout and the amino acid residues are grouped by secondary structures. Ligands are also included in the circular layout. An interaction between a pair of residues is shown as a chord connecting the two residues, and an interaction between a residue and a ligand is shown as a chord connecting the residue and the ligand. Finally, in the 3D visualization panel, the interaction patterns can be visualized on structural renderings of the proteins being compared. Both the 2D plots and 3D renderings can be downloaded by users.

### Contact Comparison Atlas: web-based analysis and visualization of interaction patterns across structures/simulations

Using GetContacts and the visualizations described above, we have built a web-based resource—the Contact Comparison Atlas (https://getcontacts.github.io/atlas/)—for analyzing interaction patterns across structures and simulations of proteins within a given family. As a proof-of-principle, we have pre-populated the Contact Comparison Atlas with interactions identified from structures/simulations of proteins in three biomedically important families: GPCRs, G proteins, and kinases. The list of structures and simulations is made available in a table and the user can select any combination of structures or simulations for comparison (Figure 3). For other proteins and protein families of interest to the user, we provide an easy-to-follow protocol for computing, comparing, and visualizing interaction patterns. Below, we describe biological insights obtained in different protein families using GetContacts and the Contact Comparison Atlas.

## Interaction patterns in GPCR structures and simulations reveal distinct and shared functional features

GPCRs are the largest family of membrane proteins in humans (~800 members) and the largest class of human drug targets. GPCRs share a conserved structural architecture with seven transmembrane α-helices (TM helices 1–7) and their activation is facilitated by extracellular ligands that favour the binding of intracellular partners such as G proteins.

### Interaction pattern in binding pockets of opioid receptors reveals conserved water-mediated interaction

Opioid receptors (subtypes: mu, delta, kappa) are a subfamily of GPCRs that play important roles in mediating central nervous system function (Stein 2016). They are also important drug targets for the treatment of pain (Matthes et al. 1996)(Vanderah 2010) and have potential for development as targets for the treatment of brain disorders (PMID: 21925742). Recent high-resolution structures of opioid receptors have revealed the presence of water molecules in the transmembrane region (Venkatakrishnan et al. 2019). Some of the water molecules are involved in mediating hydrogen bonds between the ligand and the receptor. Given the importance of water molecules for ligand binding in proteins, how similar are the water-mediated interactions in the binding-pockets of different opioid receptors?

We investigated the water-mediated interactions in opioid receptors using the Contact Comparison Atlas. Using the GPCR table there, we picked structures of opioid receptors: mu-opioid (PDB ID: 4DKL), delta-opioid (PDB ID: 4N6H) and kappa-opioid receptors (PDB ID: 4DJH). In order to eliminate differences arising due to functional state, we focused on the inactive state-structures of the three opioid receptors. After selecting these three structures, using the interaction-types panel, we examined water-mediated interactions. Interestingly, we found a conserved water-mediated interaction between the co-crystallized ligand and the receptor in all the three receptors (Figure 4a). In particular, a conserved histidine in TM6—His 6×52—forms a water-mediated interaction with a shared hydroxyl group in the ligands in all the three structures (Figure 4a). This shared hydroxyl group is conserved beyond just the small-molecule opioid ligands as the structurally equivalent interaction involving hydroxyl group of Tyr1 has also been observed in the peptide ligand bound structure of the delta opioid receptor (PDB ID: 4RWD). Thus, using the Contact Comparison Atlas, we have been able to identify a conserved water-mediated interaction across the binding-pockets of different opioid receptors. This information can be used to guide virtual screening of compounds during the hit finding phase of drug design efforts focusing on opioid receptors. Similar patterns of ligand-receptor interactions may be identified in other GPCRs subfamilies as well as across GPCRs from different subfamilies but with similar binding pockets (Lin et al. 2013).

**Fig. 4.**
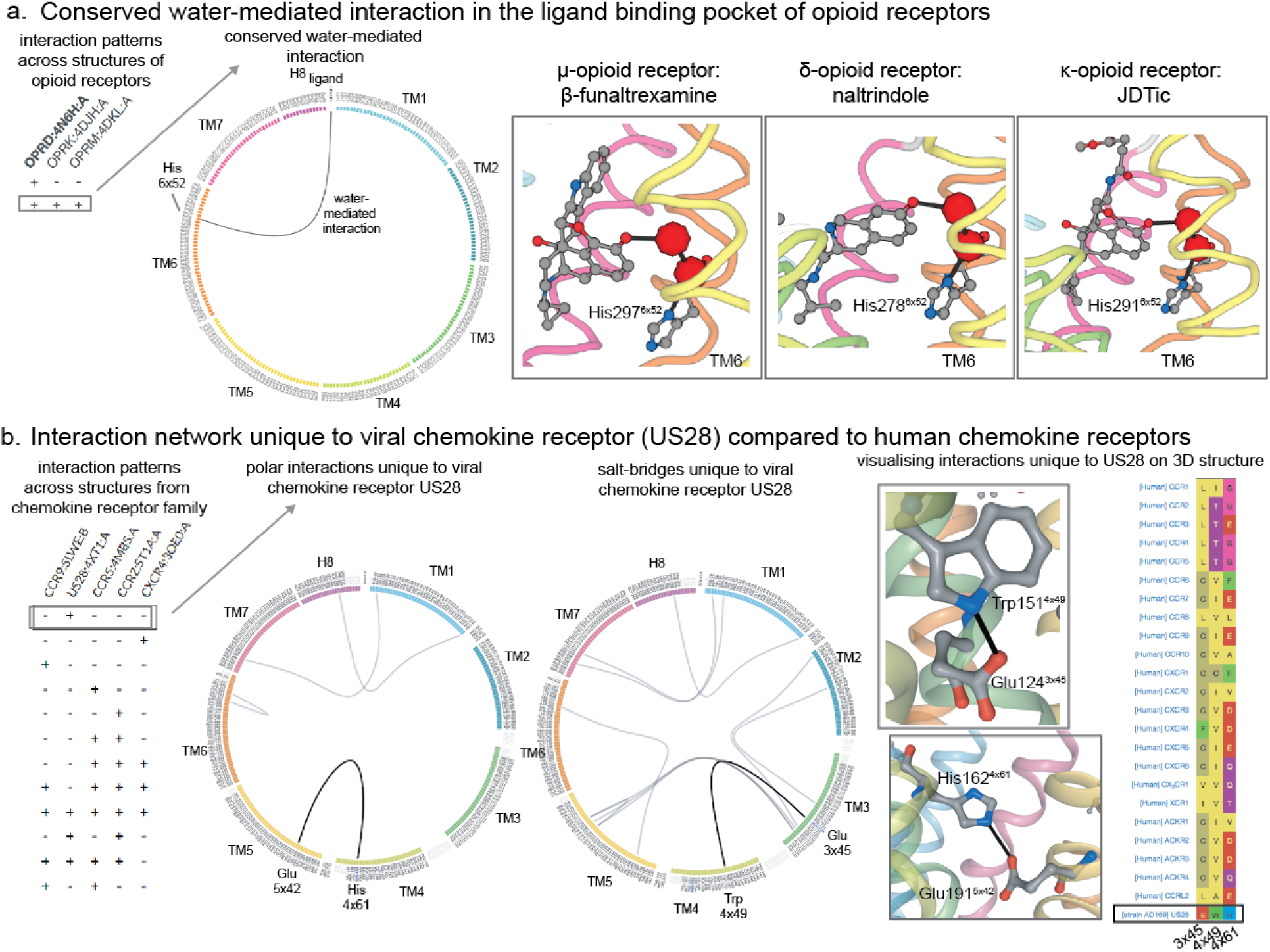
Comparison of residue interactions across structures of GPCRs. **(a)** Conserved water-mediated interaction in the ligand-binding pockets of opioid receptors. The left-hand side panel shows flareplot highlighting residue names on the circle and water-mediated interaction connecting the residues as a chord. The right-hand side panel shows the conserved water-mediated interaction between the ligand and His6×52 on TM6 of mu-, delta- and kappa-opioid receptors are shown. **(b)** Polar interaction network unique to viral chemokine receptor compared to human chemokine receptors. The left-hand side panel shows interaction patterns present across structures of the chemokine receptor family. The central panel shows flareplots displaying the network of interactions that are unique to the viral chemokine receptor US28. The right-hand side panel highlights examples of interactions that are unique to the viral chemokine receptor US28.

### Unique network of interactions in the constitutively active viral chemokine receptor US28

US28 is a viral chemokine receptor encoded by human cytomegalovirus, which is a herpesvirus that has been associated with infections in immunodeficient patients and infants (Vischer et al. 2014) and in accelerating glioblastoma growth (Heukers et al. 2018). Despite US28’s high structural similarity to human chemokine receptors, US28’s function differs from the chemokine receptors in that it shows constitutive activity. What molecular factors contribute to US28’s constitutive activity?

Using the Contact Comparison Atlas, we compared US28 with all the chemokine receptors for which structures are available: CCR2, CCR5, CCR9 and CXCR4. We found that US28 has many interactions that are distinct from the other chemokine receptors (Figure 4b). Specifically, there is a hydrogen bond between TM3 and TM4 mediated by Glu3×45 and Trp4×49. In simulations of US28, Glu3×45 was found to form an “ionic hook” with ICL2 that potentially destabilizes the inactive state (Burg et al. 2015). Furthermore, these amino acids (Glu at 3×45 and Trp at 4×45) are distinct to US28 in comparison to equivalent positions on TM3 and TM4 of the sequences of human chemokine receptors. Such interactions that are specific to US28’s structure (compared to the other chemokine receptor structures) mediated by amino acids that are distinct to the US28 sequence (compared to other chemokine receptor sequences) may be contributing to US28’s constitutive activity.

### Interaction patterns in simulations of muscarinic receptors reveal shared and distinct ligand-binding and G-protein-coupling features

Muscarinic acetylcholine receptors (M1–M5 subtypes) are important drug targets in the treatment of not only psychiatric and neurological disorders such as schizophrenia, Parkinson’s, and Alzheimer’s, but also bladder dysfunction and chronic obstructive pulmonary disease (Wess, Eglen, and Gautam 2007). All five subtypes bind the same native ligand, acetylcholine (ACh), but M2 and M4 couple to Gi whereas M1, M3 and M5 couple to Gq. How do muscarinic receptors achieve the shared ligand-binding function while maintaining differences in G-protein-coupling preference? Here, we address this question by investigating the similarities and differences in the dynamics of the muscarinic receptors.

Using GetContacts we analyzed atomic interactions from over 120 microseconds of MD simulation trajectories of the muscarinic receptors subtypes M1–4. The simulation conditions for each subtype included: (i) agonist-bound active state, (ii) unliganded active state, (iii) antagonist-bound inactive state and (iv) unliganded inactive state (Figure 5). No simulations were carried out for M5, owing to the lack of an experimental structure. We computed the interaction frequencies for all of the simulations and compared interaction frequencies between structurally equivalent pairs of residues.

**Fig. 5.**
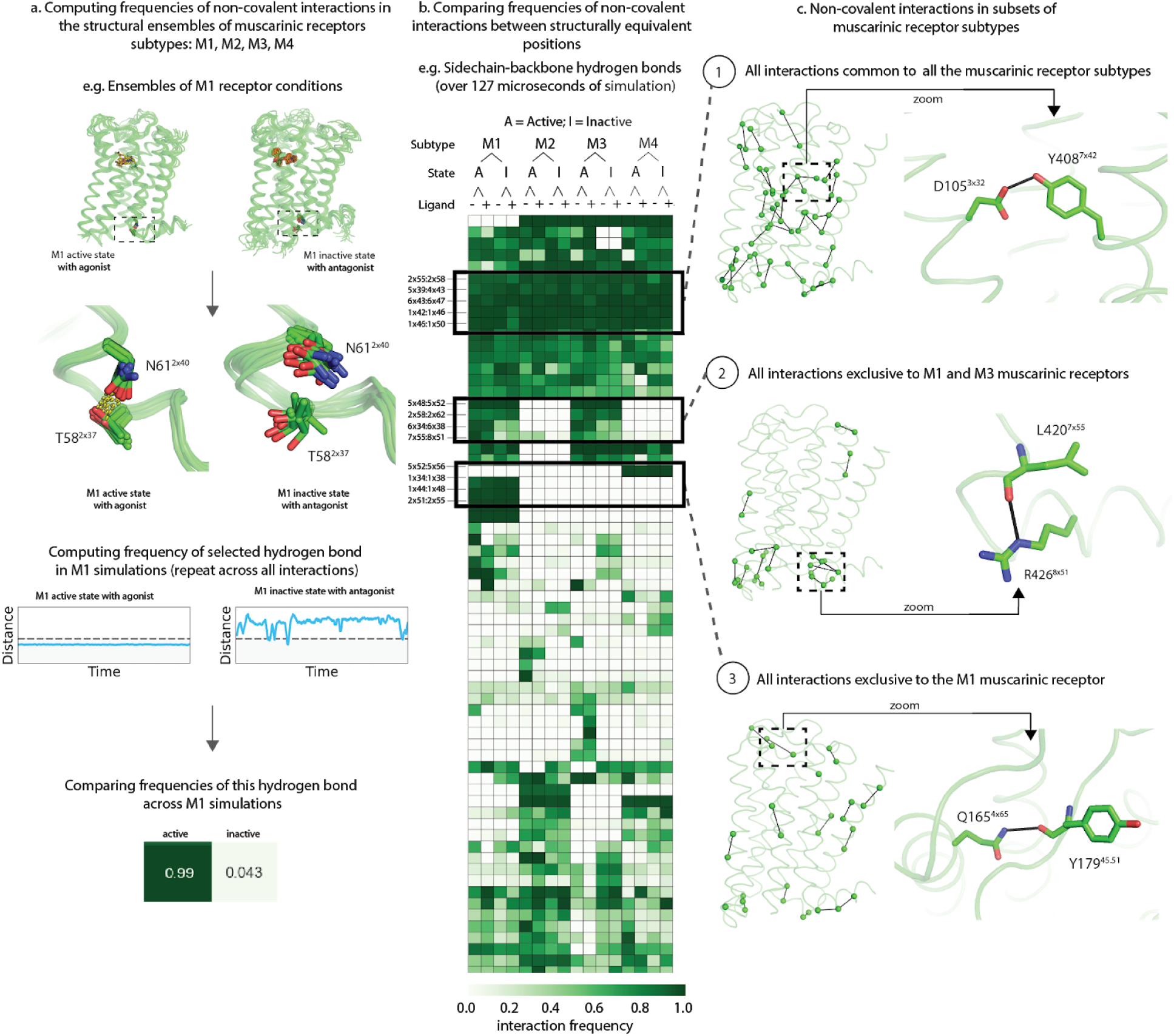
Comparison of residue interactions across simulations of muscarinic receptor subtypes. **(a)** Computing frequencies of residue interactions in simulations of the M1-4 muscarinic receptor subtypes in a variety of conditions: agonist-bound active state, unliganded active state, antagonist-bound inactive state, and unliganded inactive state, exemplified using the M1 receptor in agonist-bound active state and antagonist-bound inactive state. After the interactions observed in the trajectory are identified, an interaction frequency is computed for easy comparison across all simulation trajectories of muscarinic receptor subtypes. **(b)** Frequencies are calculated for all interactions types and filtered to include only interactions that are found to have a frequency over 50% in at least one condition. Here, one such interaction type (side chain-backbone hydrogen bonds) is displayed for simplicity. Interaction frequency is represented using a scale of white (0%) to green (100%). Interactions are further broken down by type, such as sidechain-backbone interactions (shown). Several blocks that correspond to interactions only found in unique combinations of subtypes are further highlighted in boxes. **(c)** Interaction blocks across all mAChR subtypes and interaction types that were found to be present in either all subtypes (top), M1 and M3 only (middle) or M1 only (bottom) are shown. In each case, structure of the M1 receptor is displayed in green with all interactions displayed as lines connected the C-alphas of each residue on the left, with one specific interaction highlighted on the right.

We observed that 194 hydrogen bonds were maintained with interaction frequency > 50% in at least one simulation condition (Figure 5). While the sequence similarities of M1–4 are roughly 50%, only 21.6% of the identified interactions were maintained in every subtype in at least any one of the simulation conditions. Further, only eight (or less than 0.5%) of these unique polar interactions were maintained in every simulation condition. This surprisingly small fraction of shared interactions illustrates that the majority of the identified interactions are indeed unique to some combination of muscarinic receptor subtype and condition.

Focusing on the interaction patterns provided interesting insights into shared ligand-binding properties and distinct G-protein-coupling properties (Figure 5). First, we found that interactions that are shared across all four subtypes in at least one simulation condition cluster around the orthosteric binding site (*e.g.* Asp3×32:Tyr7×42; Figure 5c, block 1). The clustering of these interactions near the binding site in all four receptors suggests a shared role of these interactions in ligand binding. Next, we identified that the interactions that are shared between M1 and M3 (which couple to Gq) but not M2 and M4 (which couple to Gi) cluster at the G-protein-binding site (Figure 5c, block 2). This clustering suggests a direct link between divergent structural interactions and function. Finally, we identified interactions that are unique to a single subtype, for example M1 (Figure 5c, block 3). These unique subtype-specific interactions could be exploited for the design of subtype-selective ligands by identifying subtype-specific conformations. For example, one of the interactions unique to M1 is a hydrogen bond between TM4 (Gln4×65) and the second extracellular loop (Tyr45×51). This interaction only occurs when Q4×65 adopts a non-crystallographic conformation to form a hydrogen bond with Tyr45×51 (Figure 5c, block 3). This new interaction changes the conformation of ECL2 and the key modulator binding residue Tyr45×51 (Abdul-Ridha, Robert Lane, et al. 2014)(Abdul-Ridha, López, et al. 2014). Different conformations of the allosteric binding site can play key roles in determining subtype selectivity, as shown in our recent study (Hollingsworth et al. 2019).

### Interaction patterns in heterotrimeric G protein structures reveals isotype-specific contacts important for regulation of the basal nucleotide exchange rate

Heterotrimeric G proteins are important transducers for GPCR-mediated signaling and are composed of three subunits, Gα, Gβ, and Gγ. In the inactive state, the Gα subunit is bound to GDP and the obligate heterodimer Gβγ, while GPCR-mediated activation of the G protein leads to nucleotide exchange of GDP for GTP (Hilger, Masureel, and Kobilka 2018). The nucleotide exchange subsequently modulates of downstream pathways through the subunits (Gα and Gβγ). In the human genome, 16 individual genes have been found to encode for 21 Gα isotypes that can be classified into four main families: G_i/o_, G_s_, G_q/11_, and G_12/13_ (Downes and Gautam 1999). It has been shown that the spontaneous GDP release rate varies among different G proteins by over two orders of magnitude (Gα_o_: 0.3 min-1, Gα_s_ 0.18 min-1, Gα_i_: 0.04 min-1, Gα_q_: 0.0016 min-1, Gα_t_: 0.00086 min-1)(Chidiac, Markin, and Ross 1999) (Ferguson et al. 1986) (Higashijima et al. 1987) (Iiri et al. 1994) (Marin et al. 2001). However, it is unclear what determines the isotype-specific spontaneous GDP release rate. It is important to understand the molecular basis of basal exchange rate as mutations in G proteins that increase this have been shown to cause different diseases including testotoxicosis and pseudohypoparathyroidism type Ia (PHP-Ia) (Markby, Onrust, and Bourne 1993) (Markby et al., 1993) and have been proposed to have oncogenic potential due to enhanced G protein signaling (Garcia-Marcos, Ghosh, and Farquhar 2011) (Garcia-Marcos et al., 2011).

The Gα subunits of heterotrimeric G proteins share a common architecture that is composed of two domains, the GTPase domain (G-domain or Ras-like domain) and the alpha helical domain (H-domain) (Oldham and Hamm 2006). The guanine nucleotide binding site is located at the interface between both domains, although most of the important contacts for GDP and GTP binding are provided by the G-domain (Sprang, 1997). Using the Contact Comparison Atlas, in the G_q_ heterotrimer structure (Nishimura et al., 2010), we identified two isotype-specific clusters of residues that form polar interactions across the domain interface. These polar interactions most likely stabilize the packing between the G- and the H-domain to prevent spontaneous dissociation of the bound nucleotide. The two clusters are formed by polar interactions of the side chains of Y291^G.hgh4.01^, R149^H.HD.11^, D236^G.s4h3.05^ and H-bonds between K57^G.H1.06^ and the backbone carbonyls of V179^H.HF.04^ and V182^G.hfs2.01^. Most interestingly, an additional stabilizing interaction specifically found in Gα_q_ involves a H-bond between the side chain of S251^G.H3.05^ in H3 and T47^G.s1h1.02^ in the P-loop, which is important for the coordination of the a- and b-phosphates of the nucleotide (Figure 6a) (Sprang, 1997). While the serine residue in H3 is highly conserved across all G protein families (except of Gα_s_), the threonine side chain in the P-loop is only conserved in G proteins of the G_q/11_ family (and G_z_) (Figure 6b). All other G protein isotypes possess an alanine at this position, which is unable to form a H-bond with residue S^G.H3.05^. In order to better understand the role of this interaction for the stabilization of the P-loop and the low basal nucleotide exchange rate of Gα_q_, we mutated the analogous alanine residue in Gα_i1_ to threonine (A41T^G.s1h1.02^) and analyzed the spontaneous GDP release. Bodipy-GTPyS binding kinetics measured by fluorescence quenching showed that insertion of this contact into Gα_i_, which shows a 25-fold faster GDP off-rate in comparison to Gα_q_, leads to a significantly reduced basal nucleotide exchange rate, under condition in which GDP release is rate-limiting (Figure 6c). This suggests that this polar contact contributes to the slow spontaneous GDP release in Gα_q_, by stabilizing the P-loop that is important for high affinity binding of the nucleotide.

**Figure 6.**
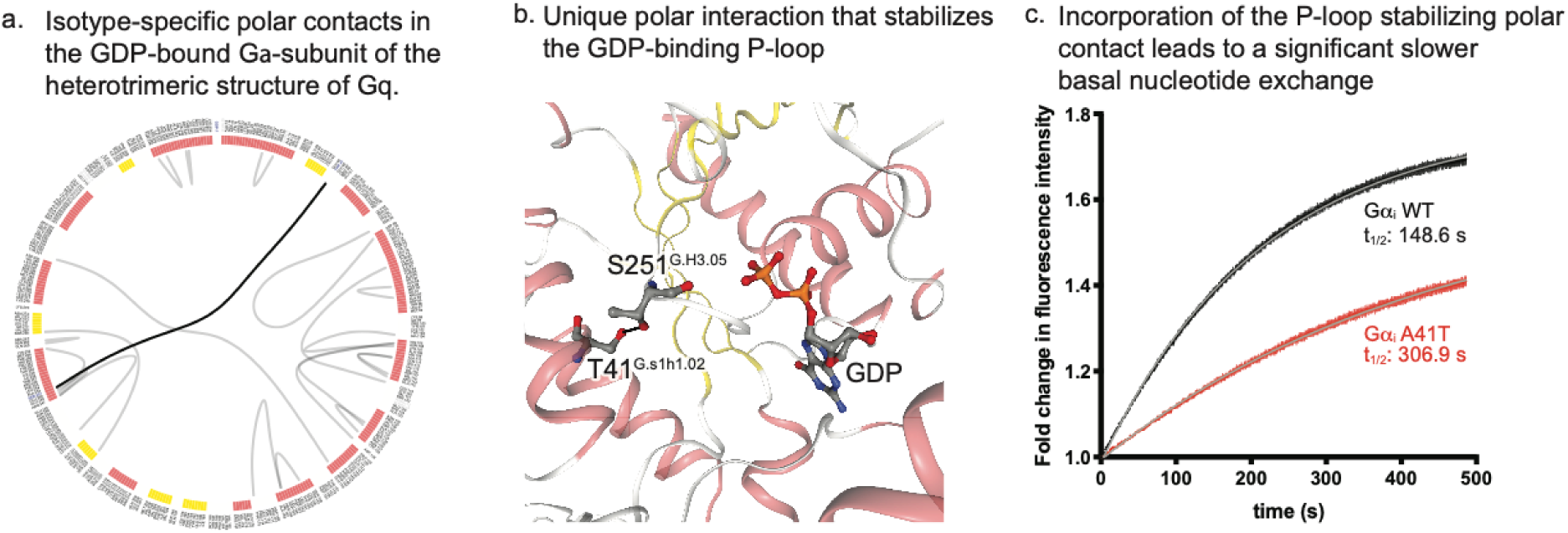
Comparison of residue interactions across heterotrimeric structure of G protein isotypes. (a) Conserved polar interactions in the Ga-subunit of heterotrimeric G protein structures. The flareplot highlights residue names on the circle and polar interactions connecting the residues as a chord. (b) Isotype-specific polar contacts in the GDP-bound Ga-subunit of the heterotrimeric structure of G_q_. The flare plot displays the network of polar interactions that are unique to the Ga_q_ subunit. The H-bond interaction between S251^G.H3.05^ in H3 and T47^G.s1h1.02^ in the P-loop is highlighted in black. (c) Unique polar interaction between S251^G.H3.05^ and T47^G.s1h1.02^ in Ga_q_ that stabilizes the GDP-binding P-loop (PDB-code 3AH8). (d) Incorporation of the P-loop stabilizing polar contact into Ga_i_ (A41T^G.s1h1.02^) leads to a significant slower basal nucleotide exchange in comparison to the Ga_i_ wild type, under conditions where GDP release is rate-limiting. GDP release was monitored by BODIPY-GTPgS binding kinetics, shown for the Ga_i_ wild type (black) and Ga_i_-A41T^G.s1h1.02^ (red). The curves represent the mean ± standard error (SEM) of three independent experiments. Fits are shown as grey lines.

## Subfamily-level comparison of kinases reveals a salt-bridge unique to tyrosine kinases

Protein kinases are the second largest protein superfamily in human with 555 members, which have been divided into nine main kinase groups (Figure 7a), such as tyrosine kinases (TK) (Manning et al. 2002). The central regulatory role of kinases in cell signaling has made them one of the most highly drugged protein families in a diversity of disease areas including cancers and neurodegenerative disorders with over 50 FDA-approved inhibitors (Carles et al. 2018). Of the eight eukaryotic protein kinase groups, the group of tyrosine kinases is the largest and the most targeted by currently approved kinase inhibitors. This group includes well-known kinases such as ABL1, EGFR and FGFR. What molecular factors contribute to making tyrosine kinases different from the other kinase groups?

**Figure 7.**
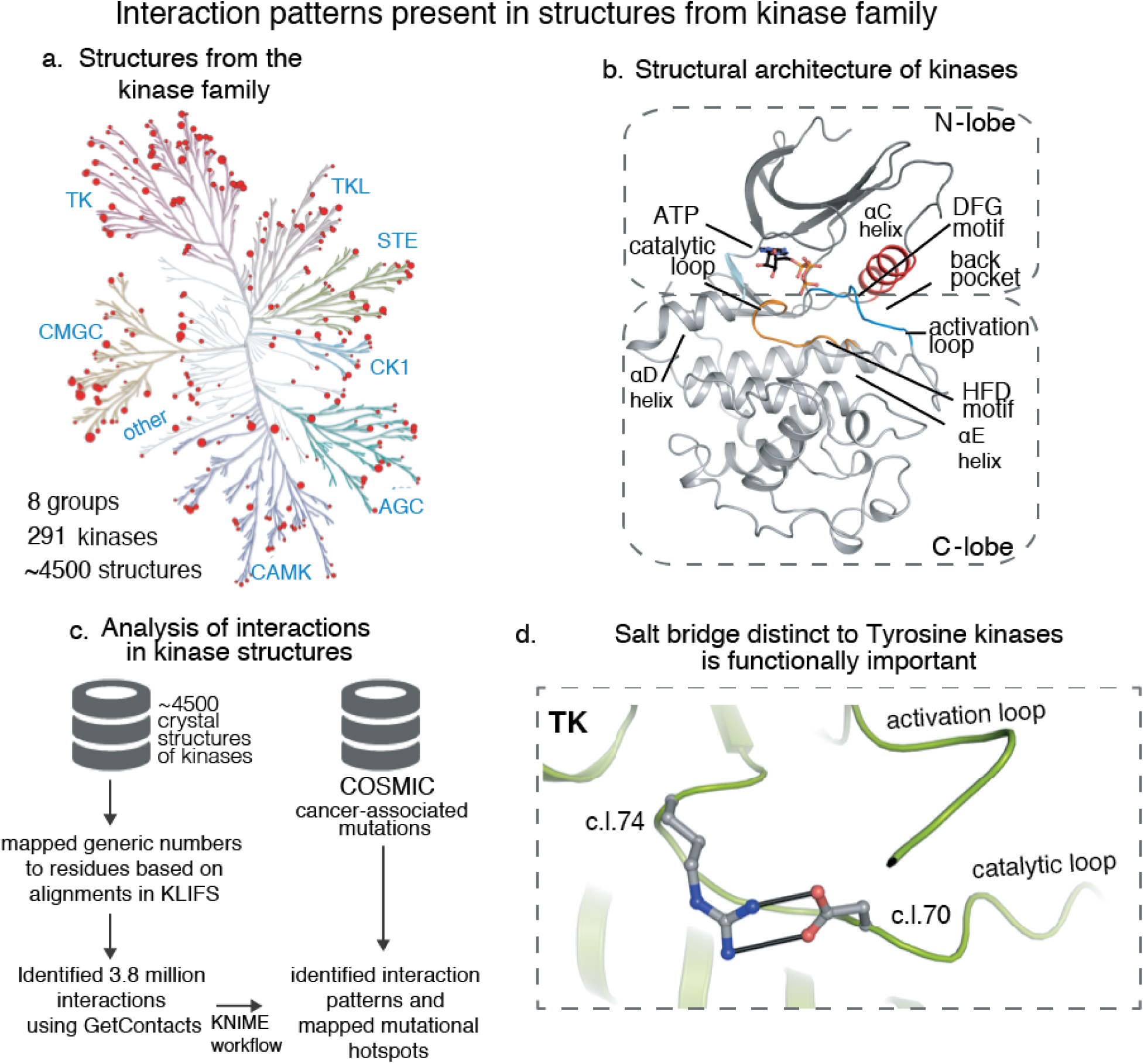
Top-down analysis of interactions in kinases: conserved interaction networks in different activation states, kinase groups, individual kinases, and binding of different inhibitors types. **(a)** An overview of the overall catalytic kinase domain. Shown is ATP in complex with CDK2 (PDB-code 1QMZ) with some characteristic kinase features highlighted: the two lobes connected by the linker region, the αC-helix, and the activation and the catalytic loop with the DFG and HRD motif respectively. **(b)** The number of available kinase crystal structures plotted onto the kinome using KinMap (the larger the dot the higher the number of available structures)(Eid et al. 2017). **(c)** A simplified workflow of the kinase interaction analysis linked to oncogenic mutations. **(d)** A conserved salt-bridge unique to the tyrosine kinase group (PDB-code 2IJM).

Structural studies of kinases have resulted in the determination of over 4500 of protein kinase crystal structures (Kooistra et al. 2016). There are 718 active-state crystal structures covering 62 out of the 95 tyrosine kinases. The availability of this vast amount of structural information provides a unique opportunity to perform comparative structural analyses towards addressing these mechanistic questions. We obtained structures of kinases from the PDB, classified them as per the KLIFS database (Kooistra et al. 2016), and used GetContacts to compute all the residue interactions for all these structures (Figure 7c). We then assigned the KLIFS numbering scheme, which captures 85 residues in the cleft between the N-lobe and C-lobe of the kinase structure.

This comparison revealed a highly-conserved salt bridge that is never observed outside the tyrosine kinase group. This salt bridge is present in about 80% of all tyrosine kinase crystal structures (Figure 7d) and is formed between the aspartate of the HRD motif (c.l.70) and an arginine residue four positions downstream (c.l.74). In cancer cell lines, mutations at this unique salt-bridge forming arginine position have been observed 95 times for kinases within the tyrosine kinase group in the COSMIC database (the median number of mutations for TKs residues was 31). Furthermore, biochemical experiments mutating this same arginine highlighted its critical function (Cai et al. 2008)(Truong et al. 2016). In one study, the mutation of the arginine to either an alanine or lysine in EGFR significantly reduced autophosphorylation activity and abolished C797 sulfenylation and redox activation (Truong et al. 2016). Another study focusing on the same arginine to lysine mutation in EGFR showed a decreased activation of diverse pathways and only a slight increase in activation of the AKT pathway compared to wild-type (Cai et al. 2008).

## Discussion and Conclusion

Studying atomic interactions in protein structures has been a useful approach, and different tools have been developed to enable computing and analyzing interactions in individual structures (Nishikawa et al. 1972; Kayikci et al. 2018). However, the true potential of interaction analysis lies in being able to perform comparisons across many structures and identify patterns that can be exploited. GetContacts and Contact Comparison Atlas build upon the strengths of previously available tools (Kayikci et al. 2018) and enable comparisons across multiple structures and simulations. They also enable efficient analysis of very large quantities of simulation and structure data. Here, we have demonstrated the use of GetContacts and Contact Comparison Atlas in identifying both shared and distinct features in various GPCRs and kinases.

A few caveats are in order. First, the interactions are identified based on the coordinates of atoms in structures and simulations. Thus, while interpreting the interactions and interaction patterns, it is recommended that users factor in the quality of the input structural data (*e.g.*, resolution) and simulation data (*e.g.*, force field accuracy and convergence of simulations). The force fields used for MD simulation are not perfect, potentially introducing artifacts in our results; simulations, however, have proven useful in elucidating a wide variety of molecular mechanisms (Hollingsworth and Dror 2018). Second, the interactions are marked as present or absent based on distance and angle cutoffs.

GetContacts and the Contact Comparison Atlas have a diversity of applications in structure modeling, drug design, protein engineering, and the interpretation of pathological mutations and genetic variation. In the context of structure modeling applications, the knowledge of interaction patterns from high-resolution structures can be used to refine homology models and identify conformational substates in simulations (Heifetz et al. 2013). In the context of drug-design applications, similarities and differences in binding-pocket interactions can be used as constraints to improve individual protein-ligand docking studies and high-throughput virtual docking screens. Furthermore, knowledge of interaction patterns across binding pockets of related targets can be useful in lead optimization for designing compounds with enhanced specificity. In the context of protein design, the knowledge of residue interactions can aid the design of novel proteins (Shen et al. 2018). For example, during the design optimization phase, the interactions that are present in lower energy design candidates can be identified. In the context of protein engineering, it is often desirable to stabilize specific conformational states. The knowledge of interactions that are distinct to a particular conformational state can be used to guide mutagenesis for structural stabilization. Finally, sequencing technologies are helping in identifying genetic variants (*e.g.* missense mutations in protein coding regions) between individuals. Comparing the residue interactions in functional regions can help with identifying disease-associated mutations from other mutations (Kayikci et al. 2018)(Hauser et al. 2018).

## Methods

### Computation of interactions from structures and simulations

GetContacts was designed and implemented to compute residue interactions in static structures and in MD simulations. GetContacts is built upon VMD-python (https://vmd.robinbetz.com/) and is capable of computing the following classes of noncovalent interactions: hydrogen bonds, salt bridges, cation-pi, pi-stacking aromatic, t-stacking aromatic, and van der Waals contacts.

Different noncovalent interactions were computed based on geometric criteria (Tina, Bhadra, and Srinivasan 2007). A salt bridge is defined to be formed between an anionic residue and a cationic residue when the distance is less than 4 Å. A cation-pi interaction is defined to be formed between a cationic residue and an aromatic residue (Phe, Tyr, and Trp) under the following conditions: the distance between cationic atom and the centroid of the aromatic ring is less than 6 Å and the angle between the vector from the aromatic centroid to the cation and the normal vector to the aromatic plane is less than 60 degrees. Pi-stacking interactions form between aromatic residues. A pi-stacking interaction is defined as formed if the distance between the aromatic centers is less than 7 Å, the angle between the normal vectors projecting from the aromatic planes is less than 30 degrees, and the angle between the first aromatic plane and the vector from the center of the first aromatic plane to the center of the second aromatic plane is less than 45 degrees. T-stacking interactions have a similar geometric criterion to Pi-stacking interactions. In this case, the distance between the aromatic centers must be less than 5 Å, the angle between the normal vectors projecting from the aromatic planes is greater than 60 degrees, and the angle between the vector from the center of the first aromatic plane to the second aromatic plane and this vector projected on the first aromatic plane is less than 45 degrees. Van der Waals contacts between two atoms are defined to exist if the distance between their centers is less than the sum of their van der Waals radii plus an epsilon value of 0.25 Å.

Hydrogen bonds are computed using the HBonds plugin in VMD (Humphrey, Dalke, and Schulten 1996), requiring the donor-to-acceptor distance to be less than 3.5 Å and the donor-hydrogen-acceptor angle larger than 110 degrees. Hydrogens are added to crystallographic structures using PyMOL (Schrödinger). We further classify hydrogen bonds between pairs of residues into three subtypes (backbone-backbone, sidechain-backbone, and sidechain-sidechain), depending on whether the interacting atoms are part of the protein backbone or of amino acid sidechains. Furthermore, we compute indirect interactions mediated through water molecules. A water-mediated interaction is defined to occur between any two atoms from a pair of residues *aa1* and *aa2* if a water molecule *w* is positioned to form hydrogen bonds *aa1*–*w* and *aa2*–*w* simultaneously. An extended water-mediated interaction forms between any two atoms in a pair of residues *aa1* and *aa2* if two water molecules *w1* and *w2* form hydrogen bonds *aa1–w1*, *w1–w2*, and *w2–aa2* simultaneously. In addition to interactions involving pairs of residues, we also compute direct hydrogen bond interactions as well as water-mediated interactions involving the ligand and residues in the binding pocket.

### Visualization of non-covalent interactions and Contact Comparison Atlas

Non-covalent contacts are visualized using flareplots (https://gpcrviz.github.io/flareplot/). The Contact Comparison Atlas was implemented primarily in JavaScript, Python, and d3.js. Flareplots have a circular layout, where the nodes represent amino acid residues as points on the circle and the interactions between the nodes are represented as edges connecting them. The interactions can be highlighted by clicking on the nodes. An interaction patterns panel is available to explore interaction patterns across different structures. Each row of this panel can be clicked and will highlight the interactions present in structures with a “+” symbol but not in those with a “−”. The accompanying interactions are also visualized on a structural rendering powered by NGLviewer (Rose et al. 2018). The structures for GPCRs, G proteins and kinases were obtained from the PDB. The interactions are computed using the GetContacts software package. The residue labels for GPCRs, G proteins and kinases were mapped using GPCRdb numbers, CGN numbers and KLIFs numbers, respectively.

### Molecular dynamics simulation system setup

Molecular dynamics simulations of the inactive states of the M1, M2, M3, and M4 mAChRs were initiated from the respective inactive-state crystal structures (M1, 5CXV; M2, 3UON; M3, 4U15; M4, 5DSG)(Haga et al. 2012; Thal et al. 2016; Thorsen et al. 2014). For inactive Apo simulations, co-crystallized ligands were removed. In inactive-state tiotropium (Tio) bound simulations, the crystallographic coordinates of Tio were used, with the exception of the M2 mAChR, for which the co-crystallized antagonist QNB was replaced with Tio (Haga et al. 2012) (for consistency, we wished to perform all inactive-state antagonist-bound simulations with the same antagonist). The experimentally determined M2 mAChR active state structure (4MQT) was used as the starting point for M2 mAChR active simulations (Kruse et al. 2013). Homology models of the active states of the M1, M3 and M4 mAChRs were constructed using Prime (Schrödinger) based on the available M2 mAChR active-state structure (4MQT) for the transmembrane region and the corresponding inactive-state structure for all extracellular and intracellular loops. ACh was modeled into the orthosteric site for all M1, M3 and M4 active-state ACh-bound conditions by first docking using Glide (Schrödinger) and then selecting a pose that replicated key ligand–receptor interactions observed in mAChR crystal structures, namely the salt bridge between the cationic choline group and D3×32 and hydrogen bonding between the acetyl group and the N6.52 side chain. The crystallographic coordinates of Ixo were used for the M2 active state Ixo-bound condition.

For each structure, we first removed any additional crystallized proteins (such as nanobodies or lysozyme) and all other non-receptor molecules with the exception of select ligands described above, and retained crystallographic waters. Prime (Schrödinger) was used to model missing side chains and loops (with the exception of ICL3, which was omitted from each receptor), and neutral acetyl and methylamide groups were added to cap protein termini. We retained titratable residues in their dominant protonation state at pH 7.0, except for residues D2×50 and D3×49, whose protonation state may change upon receptor activation (Ranganathan, Dror, and Carlsson 2014; Yao et al. 2006). These two residues were set to different protonation states in different simulations. Histidines were represented with hydrogen on the epsilon nitrogen (following manual inspection to ensure that moving the hydrogen to the delta nitrogen would not help to optimize the local hydrogen bond network).

The prepared protein structures were aligned on the transmembrane helices to the “orientation of proteins in membranes” (OPM)(Lomize et al. 2006) structure of PDB 3UON. The aligned structures were then inserted into a pre-equilibrated palmitoyl-oleoyl-phosphatidylcholine (POPC) bilayer using Dabble (http://doi.org/10.5281/zenodo.836914). Sodium and chloride ions were added to neutralize each system for a final concentration of 150 mM. Water box dimensions for each system were chosen to maintain at least an 18 Å buffer between protein images in the z direction, while bilayer dimensions were chosen to maintain at least a 35 Å buffer between proteins in the x-y plane of the membrane.

### Simulation protocols

The CHARMM36 parameter set was used for protein (including CMAP correction terms), lipid, and salt ions (Best, Mittal, et al. 2012; Best, Zhu, et al. 2012; Huang and MacKerell 2013; Klauda et al. 2010; MacKerell et al. 1998). The CHARMM TIP3P model was employed for water. Parameters for ACh were generated using the CHARMM General Force Field with the ParamChem server (Vanommeslaeghe et al. 2010; Vanommeslaeghe and MacKerell 2012; Vanommeslaeghe, Raman, and MacKerell 2012). Parameters for Tio were based on previously published parameters for *N*-methylscopolamine (Dror et al. 2013), with additional parameters supplied by ParamChem. Ligand parameters are available upon request.

Three independent simulations were performed for each condition listed in Supplementary Table 3, for a total of over 127 microseconds across all subtypes and conditions. For all active-state simulations, each prepared structure was overlaid with the experimentally determined β2-Gs complex (PDB 3SN6)(Rasmussen et al. 2011). All mAChR residues that were found to be within 5 Å of Gs in the β2-Gs complex following the overlay had a 5 kcal mol^−1^ Å^−2^ harmonic restraint to the initial position placed on the respective residue heavy atoms to ensure the receptor would remain in the active state throughout simulation. No restraints were used in inactive-state simulations.

Simulations were performed using the CUDA-enabled version of PMEMD in Amber16 on one to two graphical processing units (GPUs)(Salomon-Ferrer et al. 2013). Each system underwent a similar equilibration and minimization procedure. Systems were heated in the NVT ensemble from 0K to 100K over 12.5 picoseconds (ps), then from 100 K to 310 K over 125 ps with 10 kcal mol^−1^ Å^−2^ harmonic restraints on all non-hydrogen lipid and protein atoms. Systems were then equilibrated in the NPT ensemble at 1 bar, with a starting 5 kcal mol^−1^ Å^−2^ harmonic restraint placed on all heavy protein atoms and reduced in a stepwise fashion by 1 kcal mol^−1^ Å^−2^ every 2 nanoseconds (ns) for a total of 10 ns and then by 0.1 kcal mol^−1^ Å^−2^ every 2 nanoseconds for an additional 20 ns. Production simulations were carried out in the NPT ensemble at 310 K and 1 bar using a Langevin thermostat for temperature coupling and a Monte Carlo barostat for pressure coupling. The majority of simulations employed a timestep of 2.5 femtoseconds (fs), while others employed a time step of 4 fs with hydrogen mass repartitioning (Hopkins et al. 2015) (see Supplementary Table 1 for a list of which conditions used which approach). All bond lengths involving hydrogen atoms were constrained by SHAKE. Non-bonded interactions were cut off at 9.0 Å, while long-range electrostatic interactions were calculated using the particle mesh Ewald (PME) method with an Ewald coefficient of approximately 0.31 Å and an interpolation order of 4. The Fast Fourier Transform (FFT) grid size was chosen such that the width of a single grid cell was approximately 1 Å. Trajectory snapshots were saved every 200 ps.

### Analysis protocols for molecular dynamics simulations

The AmberTools15 CPPTRAJ package was used to reimage and center all resulting trajectories (Roe and Cheatham 2013). Simulations were visualized and analyzed using VMD (Humphrey, Dalke, and Schulten 1996). Time traces from individual simulations, such as those displayed in Fig. 5, were smoothed using a moving average with a window size of 50 ns and visualized using the PyPlot package from Matplotlib.

### Purification of Gai for nucleotide binding studies

Human Ga_i1_ subunit with an amino-terminal 8x histidine tag followed by a rhinovirus 3C protease side were expressed in Rosetta 2 (DE3) cells (EMD Millipore) using pET21a. The Ga_i1_ A41T mutant was generated by using a site-directed mutagenesis method (Liu and Naismith 2008). Cells were grown in Terrific Broth to OD600 of 0.6, and protein expression was induced by addition of 0.5 mM IPTG. After 15 h of incubation at room temperature, cells were harvested and resuspended in lysis buffer (50 mM HEPES pH 7.5, 100 mM sodium chloride, 1 mM magnesium chloride, 50 μM GDP, 5 mM beta-mercaptoethanol, 5 mM imidazole, and protease inhibitors). Cells were disrupted by sonification using a 50% duty cycle, 70% power for four times 45 s. Intact cells and cell debris were subsequently removed by centrifugation and the supernatant was incubated with Ni-NTA resin for 1.5 h at 4°C. The Ni-NTA resin was washed multiple times with lysis buffer in batch and then loaded into a wide-bore glass column, and protein was eluted with lysis buffer containing 200 mM imidazole. The eluted protein was dialyzed overnight in dialysis buffer (20 mM HEPES pH 7.5, 100 mM sodium chloride, 1 mM magnesium chloride, 10 μM GDP, 5 mM beta-mercaptoethanol, and 5 mM imidazole). The amino terminal histidine tag was cleaved by adding 1:1000 w/w 3C protease into the dialysis bag. Uncleaved protein, cleaved histidine tag, and 3C protease were subsequently removed by incubation with Ni-NTA resin for 45 min at 4°C. The resin was loaded into a wide-bore glass column and the flow-through containing the Ga subunit was collected. The protein was concentrated and run on a Superdex 200 10/300 GL column in SEC buffer (20 mM HEPES pH 7.5, 100 mM sodium chloride, 1 mM magnesium chloride, 10 μM GDP, and 100 uM TCEP).

### Nucleotide binding studies

Nucleotide binding to Ga_i1_ or Ga_i1_-A41T was followed by a change in fluorescence intensity of BODIPY-FL-GTPγS (Thermo Fisher Scientific). Fluorescence was recorded with a Horiba Fluorolog spectrofluorometer. The fluorophore was exited at 495 nm and emission was detected at 508 nm at 22°C. Slit widths were set to 0.5 nm (excitation) and 10 nm (emission). All experiments were performed in imaging buffer comprised of 20 mM HEPES, pH 7.5, 100 mM sodium chloride, 10 mM magnesium chloride, and 100 μM TCEP. Kinetics data were collected with 150 nM BODIPY-FL-GTPγS in imaging buffer in the absence of G protein for 100 s to establish the baseline fluorescence intensity. Ga subunit was added with a 1:100 dilution (1 μM final G protein concentration) and rapidly mixed in the fluorescence cuvette without halting data collection (t = 0 s). Data points were acquired every second for 600 s. The resulting kinetics spectra were plotted and fitted using GraphPad Prism 8 software.

## References

Abdul-Ridha, Alaa, Laura López, Peter Keov, David M. Thal, Shailesh N. Mistry, Patrick M. Sexton, J. Robert Lane, Meritxell Canals, and Arthur Christopoulos. 2014. “Molecular Determinants of Allosteric Modulation at the M1 Muscarinic Acetylcholine Receptor.” The Journal of Biological Chemistry 289 (9): 6067–79.

Abdul-Ridha, Alaa, J. Robert Lane, Shailesh N. Mistry, Laura López, Patrick M. Sexton, Peter J. Scammells, Arthur Christopoulos, and Meritxell Canals. 2014. “Mechanistic Insights into Allosteric Structure-Function Relationships at the M1Muscarinic Acetylcholine Receptor.” The Journal of Biological Chemistry 289 (48): 33701–11.

Ballesteros, Juan A., and Harel Weinstein. 1995. “[19] Integrated Methods for the Construction of Three-Dimensional Models and Computational Probing of Structure-Function Relations in G Protein-Coupled Receptors.” In Methods in Neurosciences, 366–428.

Best, Robert B., Jeetain Mittal, Michael Feig, and Alexander D. MacKerell Jr. 2012. “Inclusion of Many-Body Effects in the Additive CHARMM Protein CMAP Potential Results in Enhanced Cooperativity of α-Helix and β-Hairpin Formation.” Biophysical Journal 103 (5): 1045–51.

Best, Robert B., Xiao Zhu, Jihyun Shim, Pedro E. M. Lopes, Jeetain Mittal, Michael Feig, and Alexander D. Mackerell Jr. 2012. “Optimization of the Additive CHARMM All-Atom Protein Force Field Targeting Improved Sampling of the Backbone φ, ψ and Side-Chain χ(1) and χ(2) Dihedral Angles.” Journal of Chemical Theory and Computation 8 (9): 3257–73.

Burg, John S., Jessica R. Ingram, A. J. Venkatakrishnan, Kevin M. Jude, Abhiram Dukkipati, Evan N. Feinberg, Alessandro Angelini, et al. 2015. “Structural Biology. Structural Basis for Chemokine Recognition and Activation of a Viral G Protein-Coupled Receptor.” Science 347 (6226): 1113–17.

Cai, C. Q., Y. Peng, M. T. Buckley, J. Wei, F. Chen, L. Liebes, W. L. Gerald, M. R. Pincus, I. Osman, and P. Lee. 2008. “Epidermal Growth Factor Receptor Activation in Prostate Cancer by Three Novel Missense Mutations.” Oncogene 27 (22): 3201–10.

Carles, Fabrice, Stéphane Bourg, Christophe Meyer, and Pascal Bonnet. 2018. “PKIDB: A Curated, Annotated and Updated Database of Protein Kinase Inhibitors in Clinical Trials.” Molecules 23 (4). https://doi.org/10.3390/molecules23040908.

Chidiac, P., V. S. Markin, and E. M. Ross. 1999. “Kinetic Control of Guanine Nucleotide Binding to Soluble Galpha(q).” Biochemical Pharmacology 58 (1): 39–48.

Doncheva, Nadezhda T., Karsten Klein, Francisco S. Domingues, and Mario Albrecht. 2011. “Analyzing and Visualizing Residue Networks of Protein Structures.” Trends in Biochemical Sciences 36 (4): 179–82.

Downes, G. B., and N. Gautam. 1999. “The G Protein Subunit Gene Families.” Genomics 62 (3): 544–52.

Dror, Ron O., Hillary F. Green, Celine Valant, David W. Borhani, James R. Valcourt, Albert C. Pan, Daniel H. Arlow, et al. 2013. “Structural Basis for Modulation of a G-Protein-Coupled Receptor by Allosteric Drugs.” Nature 503 (7475): 295–99.

Eid, Sameh, Samo Turk, Andrea Volkamer, Friedrich Rippmann, and Simone Fulle. 2017. “KinMap: A Web-Based Tool for Interactive Navigation through Human Kinome Data.” BMC Bioinformatics 18 (1): 16.

Ferguson, K. M., T. Higashijima, M. D. Smigel, and A. G. Gilman. 1986. “The Influence of Bound GDP on the Kinetics of Guanine Nucleotide Binding to G Proteins.” The Journal of Biological Chemistry 261 (16): 7393–99.

Flock, Tilman, Charles N. J. Ravarani, Dawei Sun, A. J. Venkatakrishnan, Melis Kayikci, Christopher G. Tate, Dmitry B. Veprintsev, and M. Madan Babu. 2015. “Universal Allosteric Mechanism for Gα Activation by GPCRs” Nature 524 (7564): 173–79.

Garcia-Marcos, M., P. Ghosh, and M. G. Farquhar. 2011. “Molecular Basis of a Novel Oncogenic Mutation in GNAO1.” Oncogene 30 (23): 2691–96.

Haga, Kazuko, Andrew C. Kruse, Hidetsugu Asada, Takami Yurugi-Kobayashi, Mitsunori Shiroishi, Cheng Zhang, William I. Weis, et al. 2012. “Structure of the Human M2 Muscarinic Acetylcholine Receptor Bound to an Antagonist.” Nature 482 (7386): 547–51.

Hauser, Alexander S., Sreenivas Chavali, Ikuo Masuho, Leonie J. Jahn, Kirill A. Martemyanov, David E. Gloriam, and M. Madan Babu. 2018. “Pharmacogenomics of GPCR Drug Targets.” Cell 172 (1-2): 41–54.e19.

Heifetz, Alexander, Oliver Barker, G. Benjamin Morris, Richard J. Law, Mark Slack, and Philip C. Biggin. 2013. “Toward an Understanding of Agonist Binding to Human Orexin-1 and Orexin-2 Receptors with G-Protein-Coupled Receptor Modeling and Site-Directed Mutagenesis.” Biochemistry 52 (46): 8246–60.

Heukers, Raimond, Tian Shu Fan, Raymond H. de Wit, Jeffrey R. van Senten, Timo W. M. De Groof, Maarten P. Bebelman, Tonny Lagerweij, et al. 2018. “The Constitutive Activity of the Virally Encoded Chemokine Receptor US28 Accelerates Glioblastoma Growth.” Oncogene 37 (30): 4110–21.

Higashijima, T., K. M. Ferguson, P. C. Sternweis, M. D. Smigel, and A. G. Gilman. 1987. “Effects of Mg2+ and the Beta Gamma-Subunit Complex on the Interactions of Guanine Nucleotides with G Proteins.” The Journal of Biological Chemistry 262 (2): 762–66.

Hilger, Daniel, Matthieu Masureel, and Brian K. Kobilka. 2018. “Structure and Dynamics of GPCR Signaling Complexes.” Nature Structural & Molecular Biology 25 (1): 4–12.

Hollingsworth, Scott A., and Ron O. Dror. 2018. “Molecular Dynamics Simulation for All.” Neuron 99 (6): 1129–43.

Hollingsworth, Scott A., Brendan Kelly, Celine Valant, Jordan Arthur Michaelis, Olivia Mastromihalis, Geoff Thompson, A. J. Venkatakrishnan, et al. 2019. “Cryptic Pocket Formation Underlies Allosteric Modulator Selectivity at Muscarinic GPCRs.” Nature Communications 10 (1): 3289.

Hopkins, Chad W., Scott Le Grand, Ross C. Walker, and Adrian E. Roitberg. 2015. “Long-Time-Step Molecular Dynamics through Hydrogen Mass Repartitioning.” Journal of Chemical Theory and Computation 11 (4): 1864–74.

Huang, Jing, and Alexander D. MacKerell Jr. 2013. “CHARMM36 All-Atom Additive Protein Force Field: Validation Based on Comparison to NMR Data.” Journal of Computational Chemistry 34 (25): 2135–45.

Humphrey, W., A. Dalke, and K. Schulten. 1996. “VMD: Visual Molecular Dynamics.” Journal of Molecular Graphics 14 (1): 33–38, 27-28.

Iiri, T., P. Herzmark, J. M. Nakamoto, C. van Dop, and H. R. Bourne. 1994. “Rapid GDP Release from Gs Alpha in Patients with Gain and Loss of Endocrine Function.” Nature 371 (6493): 164–68.

Isberg, Vignir, Bas Vroling, Rob van der Kant, Kang Li, Gert Vriend, and David Gloriam. 2014. “GPCRDB: An Information System for G Protein-Coupled Receptors.” Nucleic Acids Research 42 (Database issue): D422–25.

Kayikci, Melis, A. J. Venkatakrishnan, James Scott-Brown, Charles N. J. Ravarani, Tilman Flock, and M. Madan Babu. 2018. “Visualization and Analysis of Non-Covalent Contacts Using the Protein Contacts Atlas.” Nature Structural & Molecular Biology 25 (2): 185–94.

Klauda, Jeffery B., Richard M. Venable, J. Alfredo Freites, Joseph W. O’Connor, Douglas J. Tobias, Carlos Mondragon-Ramirez, Igor Vorobyov, Alexander D. MacKerell Jr, and Richard W. Pastor. 2010. “Update of the CHARMM All-Atom Additive Force Field for Lipids: Validation on Six Lipid Types.” The Journal of Physical Chemistry. B 114 (23): 7830–43.

Kooistra, Albert J., Georgi K. Kanev, Oscar P. J. van Linden, Rob Leurs, Iwan J. P. de Esch, and Chris de Graaf. 2016. “KLIFS: A Structural Kinase-Ligand Interaction Database.” Nucleic Acids Research 44 (D1): D365–71.

Kruse, Andrew C., Aaron M. Ring, Aashish Manglik, Jianxin Hu, Kelly Hu, Katrin Eitel, Harald Hübner, et al. 2013. “Activation and Allosteric Modulation of a Muscarinic Acetylcholine Receptor.” Nature 504 (7478): 101–6.

Lin, Henry, Maria F. Sassano, Bryan L. Roth, and Brian K. Shoichet. 2013. “A Pharmacological Organization of G Protein-Coupled Receptors.” Nature Methods 10 (2): 140–46.

Liu, Huanting, and James H. Naismith. 2008. “An Efficient One-Step Site-Directed Deletion, Insertion, Single and Multiple-Site Plasmid Mutagenesis Protocol.” BMC Biotechnology 8 (December): 91.

Lomize, Mikhail A., Andrei L. Lomize, Irina D. Pogozheva, and Henry I. Mosberg. 2006. “OPM: Orientations of Proteins in Membranes Database.” Bioinformatics 22 (5): 623–25.

MacKerell, A. D., D. Bashford, M. Bellott, R. L. Dunbrack, J. D. Evanseck, M. J. Field, S. Fischer, et al. 1998. “All-Atom Empirical Potential for Molecular Modeling and Dynamics Studies of Proteins.” The Journal of Physical Chemistry. B 102 (18): 3586–3616.

Manning, G., D. B. Whyte, R. Martinez, T. Hunter, and S. Sudarsanam. 2002. “The Protein Kinase Complement of the Human Genome.” Science 298 (5600): 1912–34.

Marin, E. P., A. G. Krishna, V. Archambault, E. Simuni, W. Y. Fu, and T. P. Sakmar. 2001. “The Function of Interdomain Interactions in Controlling Nucleotide Exchange Rates in Transducin.” The Journal of Biological Chemistry 276 (26): 23873–80.

Markby, D. W., R. Onrust, and H. R. Bourne. 1993. “Separate GTP Binding and GTPase Activating Domains of a G Alpha Subunit.” Science 262 (5141): 1895–1901.

Matthes, H. W., R. Maldonado, F. Simonin, O. Valverde, S. Slowe, I. Kitchen, K. Befort, et al. 1996. “Loss of Morphine-Induced Analgesia, Reward Effect and Withdrawal Symptoms in Mice Lacking the Mu-Opioid-Receptor Gene.” Nature 383 (6603): 819–23.

Ngo, Tony, Andrey V. Ilatovskiy, Alastair G. Stewart, James L. J. Coleman, Fiona M. McRobb, R. Peter Riek, Robert M. Graham, Ruben Abagyan, Irina Kufareva, and Nicola J. Smith. 2017. “Orphan Receptor Ligand Discovery by Pickpocketing Pharmacological Neighbors.” Nature Chemical Biology 13 (2): 235–42.

Nishikawa, Ken, Tatsuo Ooi, Yoshinori Isogai, and Nobuhiko Saitô. 1972. “Tertiary Structure of Proteins. I. Representation and Computation of the Conformations.” Journal of the Physical Society of Japan 32 (5): 1331–37.

Oldham, William M., and Heidi E. Hamm. 2006. “Structural Basis of Function in Heterotrimeric G Proteins.” Quarterly Reviews of Biophysics 39 (2): 117–66.

Ranganathan, Anirudh, Ron O. Dror, and Jens Carlsson. 2014. “Insights into the Role of Asp79(2.50) in β2 Adrenergic Receptor Activation from Molecular Dynamics Simulations.” Biochemistry 53 (46): 7283–96.

Rasmussen, Søren G. F., Brian T. DeVree, Yaozhong Zou, Andrew C. Kruse, Ka Young Chung, Tong Sun Kobilka, Foon Sun Thian, et al. 2011. “Crystal Structure of the β2 Adrenergic Receptor-Gs Protein Complex.” Nature 477 (7366): 549–55.

Roe, Daniel R., and Thomas E. Cheatham. 2013. “PTRAJ and CPPTRAJ: Software for Processing and Analysis of Molecular Dynamics Trajectory Data.” Journal of Chemical Theory and Computation. https://doi.org/10.1021/ct400341p.

Rose, Alexander S., Anthony R. Bradley, Yana Valasatava, Jose M. Duarte, Andreas Prlic, and Peter W. Rose. 2018. “NGL Viewer: Web-Based Molecular Graphics for Large Complexes.” Bioinformatics 34 (21): 3755–58.

Salomon-Ferrer, Romelia, Andreas W. Götz, Duncan Poole, Scott Le Grand, and Ross C. Walker. 2013. “Routine Microsecond Molecular Dynamics Simulations with AMBER on GPUs. 2. Explicit Solvent Particle Mesh Ewald.” Journal of Chemical Theory and Computation 9 (9): 3878–88.

Shen, Hao, Jorge A. Fallas, Eric Lynch, William Sheffler, Bradley Parry, Nicholas Jannetty, Justin Decarreau, et al. 2018. “De Novo Design of Self-Assembling Helical Protein Filaments.” Science 362 (6415): 705–9.

Stein, Christoph. 2016. “Opioid Receptors.” Annual Review of Medicine 67: 433–51.

Thal, David M., Bingfa Sun, Dan Feng, Vindhya Nawaratne, Katie Leach, Christian C. Felder, Mark G. Bures, et al. 2016. “Crystal Structures of the M1 and M4 Muscarinic Acetylcholine Receptors.” Nature 531 (7594): 335–40.

Thorsen, Thor Seneca, Rachel Matt, William I. Weis, and Brian K. Kobilka. 2014. “Modified T4 Lysozyme Fusion Proteins Facilitate G Protein-Coupled Receptor Crystallogenesis.” Structure 22 (11): 1657–64.

Tina, K. G., R. Bhadra, and N. Srinivasan. 2007. “PIC: Protein Interactions Calculator.” Nucleic Acids Research 35 (Web Server issue): W473–76.

Truong, Thu H., Peter Man-Un Ung, Prakash B. Palde, Candice E. Paulsen, Avner Schlessinger, and Kate S. Carroll. 2016. “Molecular Basis for Redox Activation of Epidermal Growth Factor Receptor Kinase.” Cell Chemical Biology 23 (7): 837–48.

Vanderah, Todd W. 2010. “Delta and Kappa Opioid Receptors as Suitable Drug Targets for Pain.” The Clinical Journal of Pain 26 Suppl 10 (January): S10–15.

Vanommeslaeghe, K., E. Hatcher, C. Acharya, S. Kundu, S. Zhong, J. Shim, E. Darian, et al. 2010. “CHARMM General Force Field: A Force Field for Drug-like Molecules Compatible with the CHARMM All-Atom Additive Biological Force Fields.” Journal of Computational Chemistry 31 (4): 671–90.

Vanommeslaeghe, K., and A. D. MacKerell Jr. 2012. “Automation of the CHARMM General Force Field (CGenFF) I: Bond Perception and Atom Typing.” Journal of Chemical Information and Modeling 52 (12): 3144–54.

Vanommeslaeghe, K., E. Prabhu Raman, and A. D. MacKerell Jr. 2012. “Automation of the CHARMM General Force Field (CGenFF) II: Assignment of Bonded Parameters and Partial Atomic Charges.” Journal of Chemical Information and Modeling 52 (12): 3155–68.

Vass, Márton, Albert J. Kooistra, Dehua Yang, Raymond C. Stevens, Ming-Wei Wang, and Chris de Graaf. 2018. “Chemical Diversity in the G Protein-Coupled Receptor Superfamily.” Trends in Pharmacological Sciences. https://doi.org/10.1016/j.tips.2018.02.004.

Vass, Márton, Sabina Podlewska, Iwan J. P. de Esch, Andrzej J. Bojarski, Rob Leurs, Albert J. Kooistra, and Chris de Graaf. 2018. “Aminergic GPCR–Ligand Interactions: A Chemical and Structural Map of Receptor Mutation Data.” Journal of Medicinal Chemistry. https://doi.org/10.1021/acs.jmedchem.8b00836.

Venkatakrishnan, A. J., Xavier Deupi, Guillaume Lebon, Franziska M. Heydenreich, Tilman Flock, Tamara Miljus, Santhanam Balaji, et al. 2016. “Diverse Activation Pathways in Class A GPCRs Converge near the G-Protein-Coupling Region.” Nature 536 (7617): 484–87.

Venkatakrishnan, A. J., Xavier Deupi, Guillaume Lebon, Christopher G. Tate, Gebhard F. Schertler, and M. Madan Babu. 2013. “Molecular Signatures of G-Protein-Coupled Receptors.” Nature 494 (7436): 185–94.

Venkatakrishnan, A. J., Anthony K. Ma, Rasmus Fonseca, Naomi R. Latorraca, Brendan Kelly, Robin M. Betz, Chaitanya Asawa, Brian K. Kobilka, and Ron O. Dror. 2019. “Diverse GPCRs Exhibit Conserved Water Networks for Stabilization and Activation.” Proceedings of the National Academy of Sciences of the United States of America 116 (8): 3288–93.

Vischer, Henry F., Marco Siderius, Rob Leurs, and Martine J. Smit. 2014. “Herpesvirus-Encoded GPCRs: Neglected Players in Inflammatory and Proliferative Diseases?” Nature Reviews. Drug Discovery 13 (2): 123–39.

Wess, Jürgen, Richard M. Eglen, and Dinesh Gautam. 2007. “Muscarinic Acetylcholine Receptors: Mutant Mice Provide New Insights for Drug Development.” Nature Reviews. Drug Discovery 6 (9): 721–33.

Winn, Martyn D., Charles C. Ballard, Kevin D. Cowtan, Eleanor J. Dodson, Paul Emsley, Phil R. Evans, Ronan M. Keegan, et al. 2011. “Overview of the CCP4 Suite and Current Developments.” Acta Crystallographica. Section D, Biological Crystallography 67 (Pt 4): 235–42.

wwPDB consortium. 2018. “Protein Data Bank: The Single Global Archive for 3D Macromolecular Structure Data.” Nucleic Acids Research, October. https://doi.org/10.1093/nar/gky949.

Yao, Xiaojie, Charles Parnot, Xavier Deupi, Venkata R. P. Ratnala, Gayathri Swaminath, David Farrens, and Brian Kobilka. 2006. “Coupling Ligand Structure to Specific Conformational Switches in the beta2-Adrenoceptor.” Nature Chemical Biology 2 (8): 417–22.

Zhang, Xiuwei, Tina Perica, and Sarah A. Teichmann. 2013. “Evolution of Protein Structures and Interactions from the Perspective of Residue Contact Networks.” Current Opinion in Structural Biology 23 (6): 954–63.

